# Ecological specialization under multidimensional tradeoffs

**DOI:** 10.1101/492173

**Authors:** André Amado, Paulo R. A. Campos

## Abstract

Although tradeoffs are expected to play an essential role in shaping the diversity in a community, their effects remain relatively nebulous and notoriously difficult to assess. This is especially true when multiple tradeoffs occur simultaneously. When dealing with single tradeoffs some information can be predicted based on their curvature. Does the same happen when dealing with multiple tradeoffs? What happens if the tradeoffs have opposing curvatures? To address these issues, we develop a resource-based model that encompasses multiple tradeoffs mediated by the acquisition and processing of the resources. The model considers a spatially structured population of microbial organisms that can grow on an arbitrary number of resources, which come into the system at a constant rate and diffuse through the environment. The individuals can adopt a variety of strategies through mutation constrained by tradeoffs, which renders the model adaptive. We assess population sizes and levels of ecological specialization. We find that when multiple tradeoffs are considered the classical intuition developed for single tradeoffs does not hold. The outcome can depend significantly not only on the curvature of the tradeoffs but also on resource availability.

## 1 Introduction

Most natural environments harbor a great diversity of microbial life (Hibbing et al. 2010; Gibbons and Gilbert 2015). In those environments, microbes are typically surrounded by different strains and species with whom they compete for nutrients and space. Inside the colonies, the microbes usually express many different phenotypes in such a way that they can thrive in outcompeting and displacing their neighbors (Ghoul and Mitri 2016). The outcome of this competition and the fate of the distinct evolutionary strategies, under the constraints of tradeoffs, are greatly driven by the ecological scenarios (Ghoul and Mitri 2016; de Oliveira et al. 2018; Hoyle et al. 2008). Numerous works explore how the shape of the tradeoffs together with the ecological scenario can affect the evolutionary predictions (de Oliveira et al. 2018; Hoyle et al. 2008). Other works further suggest that even the shape of the tradeoff relationship responds to environment changes (Jessup and Bohannan 2008).

Tradeoffs are considered to have a central role in determining the patterns of species diversity in ecological communities (Kneitel and Chase 2004), including the diversity in the microbial world (Ferenci 2016). Tradeoffs constrain the range of phenotypic options that are open to organisms, and result from a number of physical and biological mechanisms (Garland 2014). On the one hand, tradeoffs curb the adaptive potential of organisms, but on the other, act as a driving mechanism for diversification in the context of environmental variation (Østman et al. 2014). Diversification implies the occurrence of ecological specialization, which in its turn were traditionally related to limited niche breadth, resulting from evolutionary tradeoffs between a species ability to exploit a wide range of resources and the effectiveness with which it uses each of these (Devictor et al. 2010). Specialization levels provide indicators of adaptive responses of populations to global changes or heterogeneity in the environment (Levins 1968; Broennimann et al. 2006; Boulangeat et al. 2012). Specialized individuals are those with limited environmental tolerance and resource use range, and are expected to be more sensitive to environmental changes than generalists, thus facing a higher risk of extinction, but on the contrary, are expected to thrive under harsh environmental conditions (Büchi and Vuilleumier 2014). In fact, ecological specialization is disclosed at different biological levels: population, species, community, but also at the individual level (Devictor et al. 2010; Bolnick et al. 2002).

Despite their importance to life-history theory, quantifying tradeoffs requires longterm observation and modeling, and are subject to several limitations (Pease and Bull 1988). This problem is even more subtle when multidimensional tradeoffs exist (Edwards, Klausmeier, and Litchman 2011; Lancaster, Hazard, Clobert, and Sinervo 2008). The existence and interplay of multiple tradeoffs and their consequences to the degree of ecological specialization are addressed here. The tradeoffs are here modelled as surfaces and curves in trait space, thus representing the constraints between the variables. Our approach is developed within a resource-based modelling framework, whereby we are concerned with the metabolic machinery of the individuals. In particular, two well-acknowledged tradeoffs in the literature are considered: tradeoffs between resource uptake rates and the rate-yield tradeoff (Litchman et al. 2015). The latter one defines a constraint between the rate at which a given resource is uptaken and the efficiency of the process of conversion of the seized resources into ATP (energy). The curvature of the surface and curves in trait space can be tuned, thus allowing us to cover a broad spectrum of possible shapes. As aforesaid, this matter is of great relevance especially in the face of our knowledge from classical ecological models that deal with two-traits relationships (Egas et al. 2004; Hoyle et al. 2008). These classical ecological models state that specialization is expected under convex curves, whereas the generalist strategy is favored under concave relationships. So, many questions arise upon the occurrence of multivariate tradeoff patterns, especially those involving different classes of tradeoffs (distinct biological quantities). For instance, what is the evolutionary outcome when tradeoffs of opposing curvatures exist? And what about multidimensional tradeoffs involving different quantities? Do we have any kind of hierarchy regarding the tradeoffs? These are the types of questions we address in the current contribution.

The fact that a resource-based modelling approach is adopted, allows us to go further and investigate whether the tradeoff shape by its own is on the ground of the ultimate composition of ecological systems or ecological responses due to the existence of trade-offs also rely on environmental conditions. With these objectives in mind, we propose a structured model in which population is distributed over a two-dimensional lattice, and resources diffuse over lattice following the standard dynamics provided by a discretized version of the diffusion equation. We investigate population sizes, specialization levels and the spatial distribution of trait-values under different scenarios, and for many different shapes and combinations of tradeoffs. Specialization levels are defined at the individual level (Bolnick, Yang, Fordyce, Davis, and Svanbäck 2002), and so for each pairwise tradeoff relationship, we examine if one of the traits is overexpressed relative to the other one. Efforts are also undertaken to understand the role of structuring in the emerging patterns of specialization. As we will see, quite intricate patterns for the specialization levels emerge. The resulting patterns can not be explained by considering ecological systems subject to multivariate tradeoffs as a superposition of the predictions derived from classical ecological models of two-traits. We find that structuring also plays a central role determining the evolutionary outcomes of the system.

The paper is organized as follows. In section 2, we describe the model used to address the aforedescribed problem and the technical aspects that underlie it. This is followed by a description of the results in section 3, and the presentation of our concluding remarks in section 4. Some of our simulation results are presented as Supplemental Informations. In the same section, a summary of all parameters defined in the modelling is introduced.

## 2 Materials and Methods

Let us start by providing a brief overview of the model. The present work is developed within the framework of a resource-based modelling. We deal with a spatially structured population that acquires resources from the environment. Those resources are provided to the system at a constant rate and then diffuse over the environment, ultimately being captured by the individuals. The individuals reproduce and mutate and thus selection will optimize their resource uptake and resource processing rates. However, these rates are subject to tradeoffs which prevent their simultaneous optimization. Two types of resource related constraints are considered. First, the tradeoff between resource uptake rates, meaning that the organism cannot optimize simultaneously the uptake of all resources. This tradeoff stems from several factors, including limitation of energy for active transport of substances and the limited area of the cell, or in a different perspective the time the organism has available to acquire food. Second, the tradeoff between the uptake rate of each resource and the yield of the conversion of that resource into energy. This second tradeoff can be ultimately traced to fundamental limitations imposed by thermodynamics (Novak et al. 2006; MacLean 2008). Both types of tradeoffs are well established and documented in the literature (Pfeiffer et al. 2001; Litchman et al. 2015). Usually these tradeoffs are studied separately, but in real biological systems they act simultaneously.

The present contribution aims to comprehend how this entanglement of multiple tradeoff relationships affects the patterns of the ecological specialization. With this in mind, one assumes that a total number of *N* different types of resources are available to the individuals. The population is spatially structured, and thereby the individuals are arranged over a two-dimensional lattice of linear size *L*. At most a single individual can lie in each site of the lattice, and so each site can be either occupied or vacant. The *N* resources flow into the lattice through *n*_loc_ distinct sites. These sites are independent for each resource and are randomly chosen in the lattice for each configuration. To ensure the results for different *N* are comparable we fix the total flux of resources entering the system (lattice) as Δ*R*. This way, for each resource type, we have an inflow of Δ*R/N* units of resource per time step. After entering the lattice, the resources diffuse over it, following a discretized version of the diffusion equation

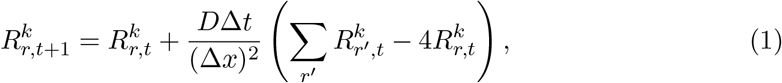

where *D* is a diffusion constant, Δ*t* is the time step, Δ*x* is the lattice spacing,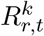 denotes the amount of resource *k* at position *r* at time *t* and the sum in *r*′ goes over the 4 adjacent neighbors. This discretized version of the diffusion equation requires the condition 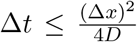 to be respected in order to avoid stability issues. As usual with diffusion, the flow of a given resource will be largest in the direction of the largest gradient.

An individual *i* is characterized by its uptake rates 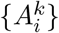, with *k* = 1*, …, N*, and their respective returns 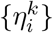, which is the yield of the process that converts biochemical energy from nutrients, here taken as a direct measure of growth rate. The biochemical energy obtained from nutrients, such as glucose, is typically used to produce adenosine triphosphate (ATP), which fuels the biochemical processes inside the cell.

Individuals can only grab resources at their locations. At each time step Δ*t*, each individual *i* at the position *r* of the lattice will seize an amount of the resource *k*, which equals

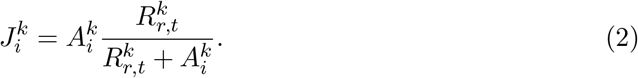

The amount of resource is calculated for each resource type. The above equation is based on Monod and Michaelis-Menten formulations (Rockwood 2015). Once the resources are captured by the individual, they will subsequently be converted into energy. Note that not all resource types are equally processed by the individual, as each one involves a different metabolic machinery, with their own constraints. So, it sounds natural to associate with each resource *k* a return *η*^*k*^, which will tell us the amount of energy extracted from that resource amount. Borrowing ideas from the microbial world, the amount of energy due to resource *k* will be

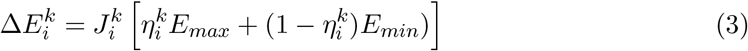

with the total change of internal energy taken as the sum over all resource types

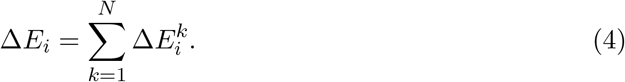

*E*_max_ is attained when one has maximum efficiency, *η*^*k*^ = 1, whereas *E*_min_ corresponds to a lower bound, when *η*^*k*^ = 0. For simplicity, we adopt as reference values *E*_max_ = 36 and *E*_min_ = 2, inspired by the metabolic machinery of single-celled organisms, such as bacteria and yeast, where the inneficient mode of metabolism (fermentation) yields typically 2 ATP molecules per glucose molecule, whereas the efficient mode of metabolism, dubbed cellular respiration, yields up to 36 ATP molecules for each glucose molecule (Campbell and Farrell 2006).

### 2.1 Biological life cycle

The following phase concerns the biological cycle, in which the individuals first seize a fraction of the resources which are available in their local site (Eq. (2)), which in turn will result in an increase in its energy reservoir (biomass). Reproduction will take place every time the individual’s energy level reaches a threshold value *E*_split_. During cell division, the parental cell divides into two cells with half of its energy level each. One of the cells remains in the focal position, whereas the second daughter cell will occupy an unoccupied site in the neighborhood of the focal cell. If there is no empty site in its neighborhood, the daughter cell will replace a random neighbor. The daughter cells are not metabolically identical to the parental cell, but present slightly different values from the parental cell due to a small mutation introduced in reproduction. In this sense, our model is adaptive, as the individuals with better phenotypic responses to the local environment are more likely to thrive, as they can better exploit the available resources and use their metabolic machinery to directly increase the reproductive rate, hence locally spreading.

The final stage regards the process of cell death, in which individuals die at a constant rate *ν*.

### 2.2 The tradeoff relationships

In the current work, we are concerned with tradeoff relationships affecting metabolic properties of the individuals. Within the perspective of resource handling, two types of tradeoffs are acknowledged, those establishing constraints between resource uptake rates, *A*^*j*^ × *A*^*k*^, and for each resource type a tradeoff between the resource uptake rate and its corresponding return, *A*^*j*^ × *η*^*j*^. This set of pairwise relations altogether results in a multidimensional coupling as the change in one of those variables leads to changes of all other quantities. A simpler and more elegant formulation is to define a surface in trait space, thus representing the allowed region of phenotypic values stemming from physical and biological constraints. Through the mutation process during reproduction, different phenotypic responses are verified and those that are locally and instantaneously more advantageous can become established.

An individual can be represented by a vector in the trait space, 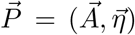, with 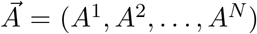 and 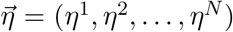, hence giving origin to a 2*N*-dimensional space. We will assume that only a subspace of this is accessible due to the tradeoff relationships and that the accessible subspace lies on a surface that defines those tradeoffs.

We intend then to explore the way the curvature of that surface shapes the specialization of the individuals at equilibrium.

To parametrize the tradeoff surface, we will make use of an adaptation of a family of polynomial functions, known as Bézier functions (Farin et al. 2002; Chen and Wang 2003). In *N* dimensions, a point in the surface is parametrized by

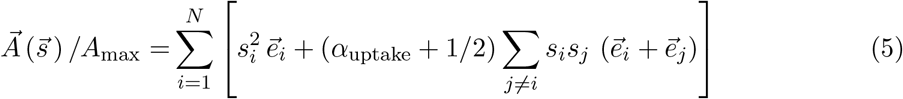

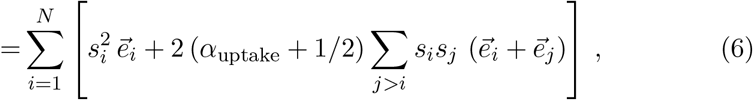

where 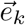 is a unit vector in the *k*-direction, with *k* corresponding to a resource index, and the condition 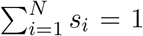 should be respected. Therefore, to fully parametrize the system *n-*1 variables are necessary. The full set of unit vectors 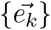 defines a complete orthonormal basis in the subspace 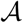. The parameter *α*_uptake_ determines the shape of the surface. In terms of its components, (*A*^1^, *A*^2^, …, *A*^*N*^), the above equation can be rewritten as

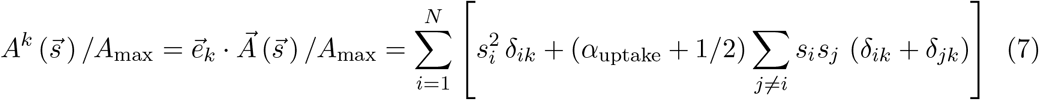

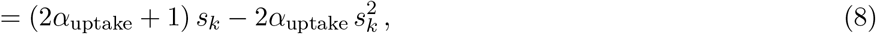

which facilitates its numerical implementation. In the above equation, *δ*_*ij*_ stands for the Kronecker delta which is 1 if *i* = *j* and 0 otherwise.

In order to determine the efficiency *η*^*k*^ associated with resource of type *k*, we assume a similar dependence to those between the uptake rates, {*A*^*k*^}, but now between *A*^*k*^ and *η*^*k*^. As this relationship is between two variables only, we can solve it explicitly, without needing to rely on *s* parameters. Therefore, the efficiency *η*^*k*^ is determined by the equation

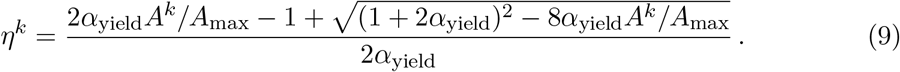

Since *A*^*k*^ can take values between zero and *A*_max_, we have to normalize it. As expected, in the limit *α*_yield_ *→* 0, which corresponds to a linear dependence, the above equation reduces to *η*^*k*^ = 1 − *A*^*k*^/*A*_max_.

As previously discussed, connected with the reproduction a small mutation is introduced. This step is implemented by randomly picking a single parameter *s*_*k*_ and modifying it by an amount *δs*_*k*_, which is a random gaussian distributed variable with mean zero and standard deviation *σ* = 0.01. The remaining parameters are then readjusted so that the condition ∑_*k*_ *s*_*k*_ = 1 continues being respected. The following step is to recalculate the resource uptake rates from the set of {*s*_*k*_}-values, and their corresponding efficiencies *η*^*k*^ are promptly evaluated from Eq. (9).

The parameters *α*_uptake_ and *α*_yield_ play a key role as they define the shape of the tradeoff relationships. Negative values of *α* generate convex surfaces, whereas positive values lead to concave surfaces. When *α* = 0, the variables are linked by linear relations. Figures 1 and 2 show intances of the surfaces produced through the Bézier functions. We can see that the *α* parameters provide an effective parametrization for the curvature of the surface, with the possibility to change from a negative to a positive curvature by tuning one parameter only, even in the multidimentional case. In the plots, the resource uptake rates are rescaled by their maximum value, *A*^*k*^/*A*_max_.

**Figure 1:**
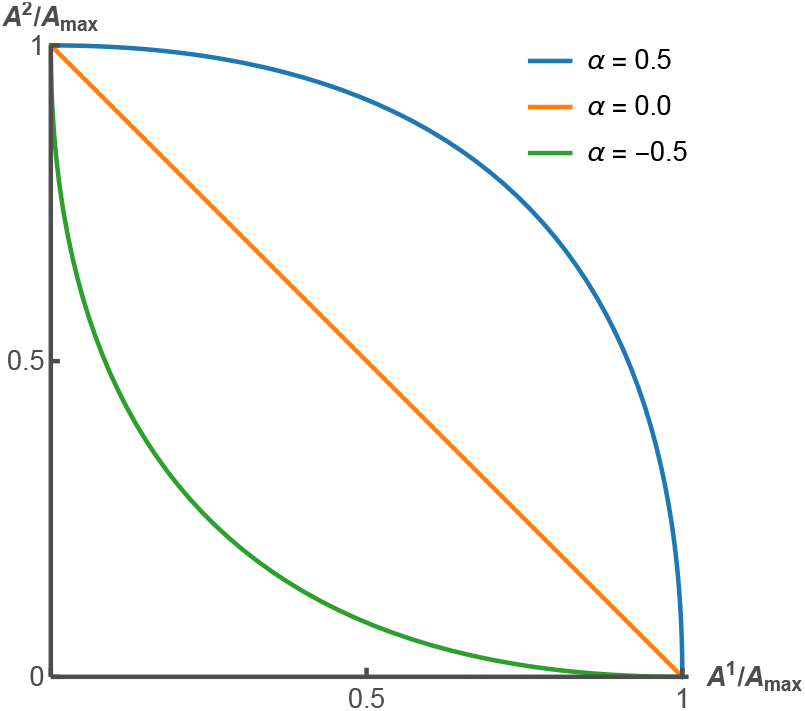
A model of two-resources. Tradeoff between *A*^1^ and *A*^2^ for *α* = −0.5 (green curve), *α* = 0 (orange curve), *α* = 0.5 (blue curve).

**Figure 2:**
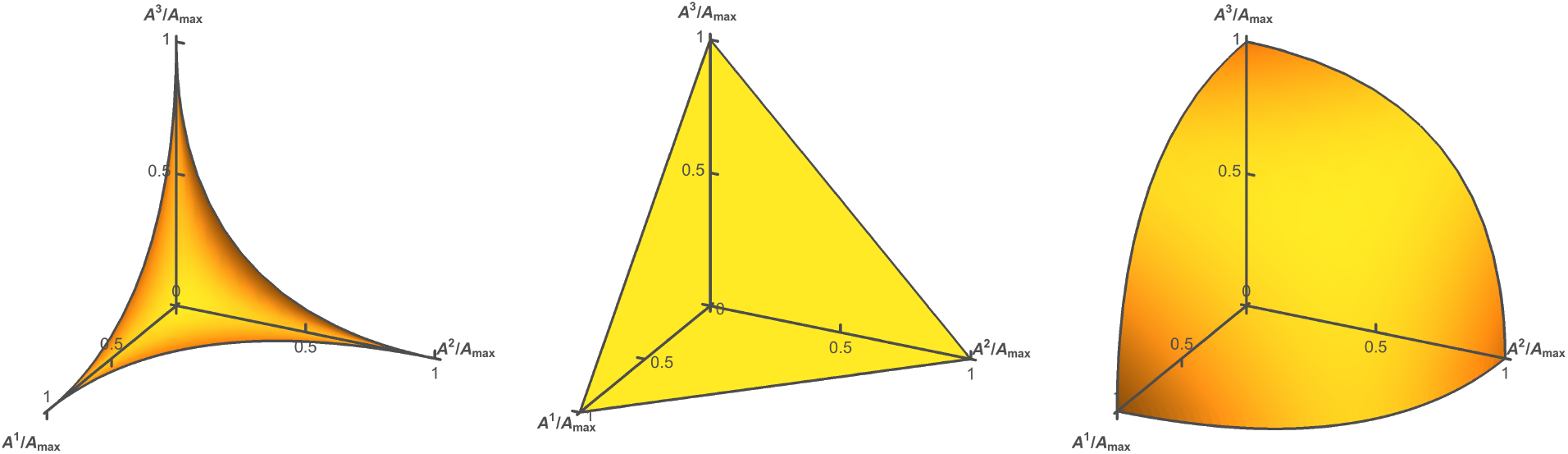
A model of three-resources. Tradeoff between *A*^1^, *A*^2^ and *A*^3^ for *α* = −0.5 (left panel), *α* = 0 (middle panel), *α* = 0.5 (right panel).

### 2.3 Specialization

Our main interest is measuring the specialization level of the individuals in the population. For that purpose it is necessary to introduce a measure of specialization. Since the phenotypic characteristics we are interested in are the uptake rate and yield of the metabolism we adopt a criterion that identifies specialization in these quantities. Hence, for the specialization in uptake, we verify if the ratio *A*^*j*^/*A*^*k*^ conforms either to *A*^*j*^/*A*^*k*^ > *r*_*spec*_ or *A*^*j*^/*A*^*k*^ < 1*/r*_*spec*_, meaning that one uptake is much larger than the other. If one of these conditions is obeyed, we consider that the individual is specialized in the uptake of resource *j* over *k* or vice versa. Therefore, each pairwise relation *A*^*j*^ × *A*^*k*^ contributes to the total specialization in the resource uptake with 0 or 1. We then sum over the different pairwise relationships and average over individuals to obtain the total specialization in resource uptake of the population. This way, in our metric the specialization level is defined in the range (0, *M*_uptake_), where *M*_uptake_ = *N ×* (*N* − 1)/2, the number of pairwise relations of the class *A*^*j*^ × *A*^*k*^. The maximum specialization *M*_uptake_ occurs whenever each of the *M*_uptake_ relationships results in the specialization in either resource for all individuals. Of course, due to conflicting relations between those variables, this maximum level may never be attained. A similar analysis is carried out for the levels of specialization in high yield, but as there are *N* pairwise relationships of the class *A*^*k*^ × *η*^*k*^, the maximum possible value of specialization in high yield is *N*. The measurement only accounts for the condition *η*^*k*^/(*A*^*k*^/*A*_*max*_) > *r*_*spec*_, implying it is directional, in the sense only those individuals that specialize in finding a high efficiency in a given resource or given set of resources contribute to the specialization in high yield.

## 3 Results

In this section we present the simulation results, aiming to determine the levels of ecological specialization under quite distinct scenarios. In our simulations, unless stated otherwise, the linear system size is *L* = 100, in such a way that there are *L× L* sites. The number of resource input locations for each resource type is *n*_loc_ = 100 and the diffusion constant *D* is set at *D* = 0.2. The measurements are taken after the system has reached an equilibrium regime.

First, it is important to determine how the population size is influenced by the variables of the model. In the left panel of Fig. 3, the dependence of the population size, here expressed as the fraction of occupied sites of the lattice, on the curvature parameter *α*_uptake_ is explored. Meanwhile, *α*_yield_ is set at *α*_yield_ = 0, meaning a linear relationship between resource uptake rate *A*^*k*^ and efficiency *η*^*k*^ for each resource type. Different values of resource input Δ*R* are simulated. As expected, the population size is an increasing function of Δ*R*. In fact, the lattice becomes almost fully populated when Δ*R* is large and *α*_uptake_ takes very negative values. Under these circumstances, space limitation is expected to play an important role. In a nutshell, one may also conclude that the occupation fraction decreases slightly with the curvature parameter *α*_uptake_. If looked at isolatedly, one would say that negative values of *α*_uptake_ favor the rise of specialized individuals in the resource uptake of one or a few resource types. Under this perspective, those results seem to demonstrate that larger populations can be accommodated when ecological specialization emerges as it allows a partitioned usage of the resources. On the other hand, we observe that by holding *α*_uptake_ constant and varying *α*_yield_ (right panel of Fig. 3), the variation of the lattice occupation with *α*_yield_ is less evident, but still, at least for large Δ*R*, the occupation fraction also decreases slightly when *α*_yield_ rises. Figs. S2-4 (Supplemental Information) present a more complete picture of how the population size changes with *α*-parameters. The figures disclose the lattice occupation for many combinations of values of *α*_uptake_ and *α*_yield_. Regardless the number of resources *N*, the scenarios are qualitatively similar. The dependence of the lattice occupation on *α*_yield_ is clearly determined by the resource influx Δ*R*. While for large Δ*R*, the lattice occupation declines as *α*_yield_ increases, a conflicting scenario emerges for small Δ*R*, as now the lattice occupation grows with *α*yield.

**Figure 3:**
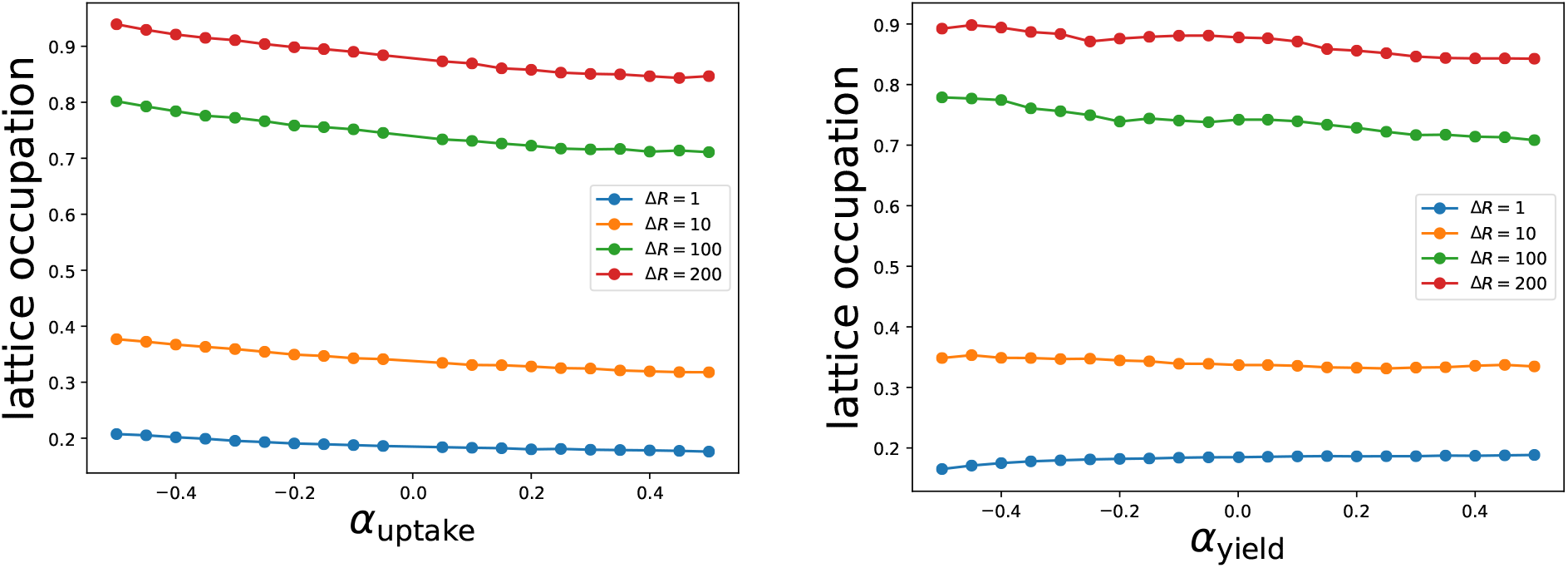
Lattice occupation as a fraction of the maximum lattice capacity for several values of resource input rate Δ*R*. The left panel depicts the dependence with *α*_uptake_ for constant *α*_yield_ = 0 and the right panel shows the dependence with *α*_yield_ for constant *α*_uptake_ = 0. The parameters are *N* = 3, *ν* = 0.01, *A*_max_ = 10, *L* = 100, *D* = 0.2, Δ*t* = 1 and Δ*x* = 1. Each point consists of an average over 100000 time units, sampled every 50 time units, taken from 50 independent configurations.

In order to check how the above results are related to the emerging specialization levels, the left panel of Fig. 4 shows the level of specialization in resource uptake against the curvature parameter *α*_uptake_. Once again, one has *N* = 3 resources and *α*_yield_ = 0. From the plot, we find out that the level of specialization is nearly a monotonic decreasing function of *α*_uptake_, allowing us to relate increased population sizes with higher levels of specialization in the uptake of resources. Note also that the variation of *α*_uptake_ also influences specialization levels in yield.

**Figure 4:**
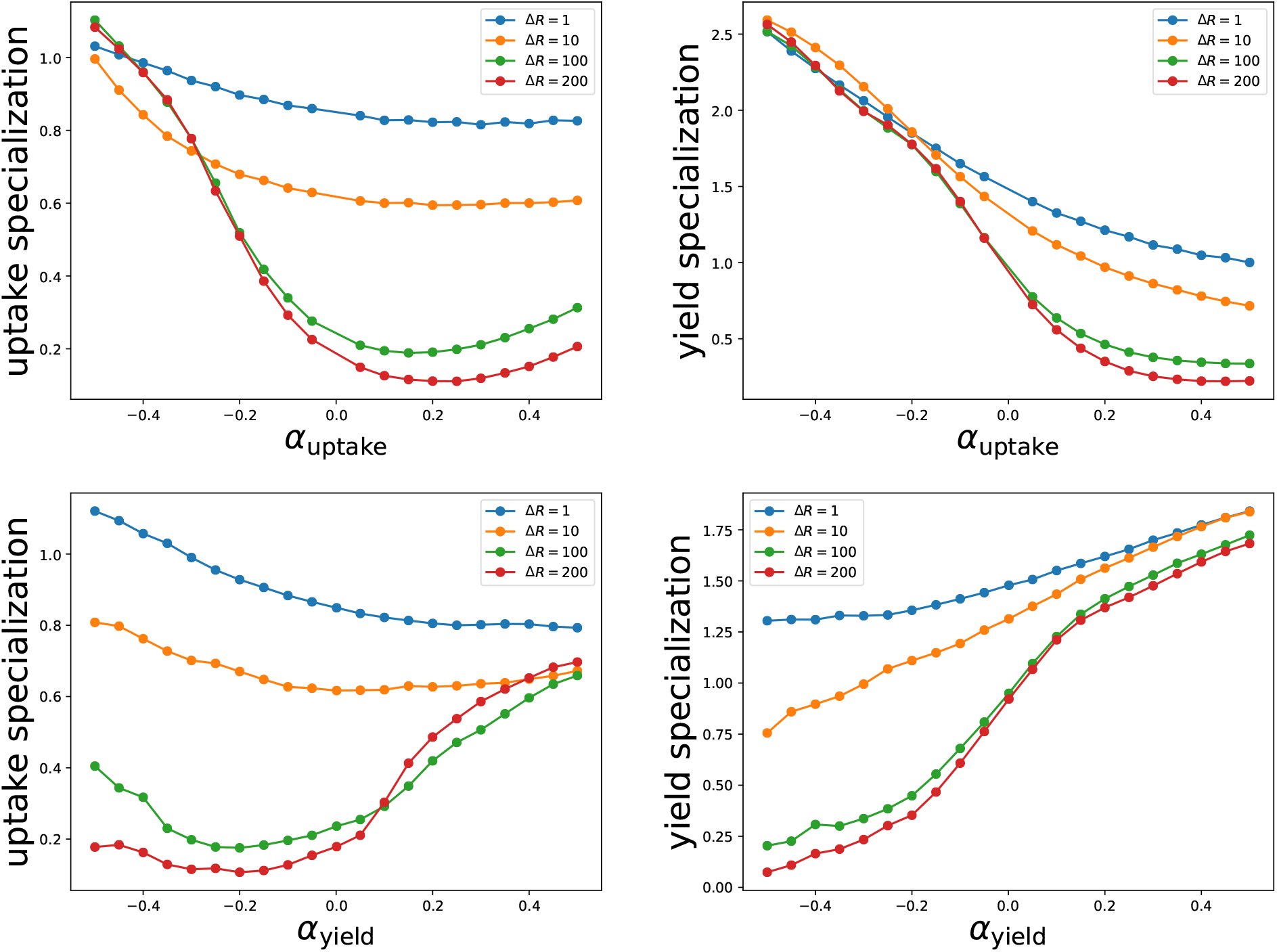
Specialization level for several values of resource input rate Δ*R*. The left panels depict the specialization level in uptake and the right panels the specialization in yield. In the top panels, *α*_uptake_ varies while *α*_yield_ varies while *α*_uptake_ is fixed at zero. The parameters are *N* = 3, *ν* = 0.01, *A*_max_ = 10, *L* = 100, *D* = 0.2, Δ*t* = 1 and Δ*x* = 1. Each point consists of an average over 100000 time units, sampled every 50 time units, taken from 50 independent configurations.

Let us now address the reversed situation, where *α*_uptake_ = 0 and *α*_yield_ is varied. Curiously, one notices that augment of *α*_yield_ causes enhanced specialization in high yield. Contrary to common sense, if lonely examined and employing the reasoning of classical ecological models of single tradeoffs, it would be expected a loss of specialization with increased *α*_yield_. Note that the growth of specialization levels in high yield is accompanied by the decline of specialization levels in resource uptake when Δ*R* has low or moderate values, while it is accompanied by the increase of specialization levels in resource uptake when Δ*R* has large values. Altogether, the results seem to propose that population sizes are more responsive to specialization in resource uptake. The simulation results in Fig. 4 also substantiate previous findings that claim that the scarcity of resources creates pressure for the population to specialize in the use of different sets of resources (Østman, Lin, and Adami 2014).

To confirm that the model yields the expected results in the case of a single tradeoff we also ran a variation of the model where the tradeoff between the rate *A*^*k*^ and the efficiency *η*^*k*^ was removed (outcomes in Fig. S1).

For the sake of simplicity, from now on the levels of specialization will be normalized by their maximum possible value, so that one can more directly compare the role of the tradeoffs under scenarios of distinct number of resources. Figure 5 presents heat maps for the specialization levels in both resource uptake and high yield for number of resources *N* = 2, and over a broad domain of the curvature parameters, *α*_uptake_ and *α*_yield_. In the upper panels one considers a resource supply rate Δ*R* = 1, whereas in the lower panels Δ*R* = 200. In the domain of positive values of *α*_yield_, the augment of *α*_uptake_ at fixed *α*_yield_ results in lower levels of specialization in both resource uptake and high yield, which is in accordance with the outcomes shown in Fig. 4. If only discussed from the perspective of the relationship *A*^1^ × *A*^2^, this behavior sounds intuitive, as concave curvatures tend to promote the generalist behavior. However, a more complicated picture comes up in the negative domain of *α*_yield_. In that range, the increase of *α*_uptake_ boosts specialization in resource uptake. Except when *α*_uptake_ is nearly −0.5, one observes a tiny interval in which specialization levels in resource uptake first drops, hence reaching a domain in which specialization in resource uptake is essentially missing before growing with a further increase of *α*_uptake_. Thus, there is a loss of generality of the outcomes posed in Figure 4. In this aspect, the issue looks much simpler whether the dependence of specialization levels in high yield on *α*_uptake_ is checked out (right panel). In all the domain, the rise of *α*_uptake_ substantially diminishes the levels of specialization in high yield, as seen when the resources are abundant (large Δ*R*), where over an extensive region of the diagram no specialization in high yield evolves at all.

**Figure 5:**
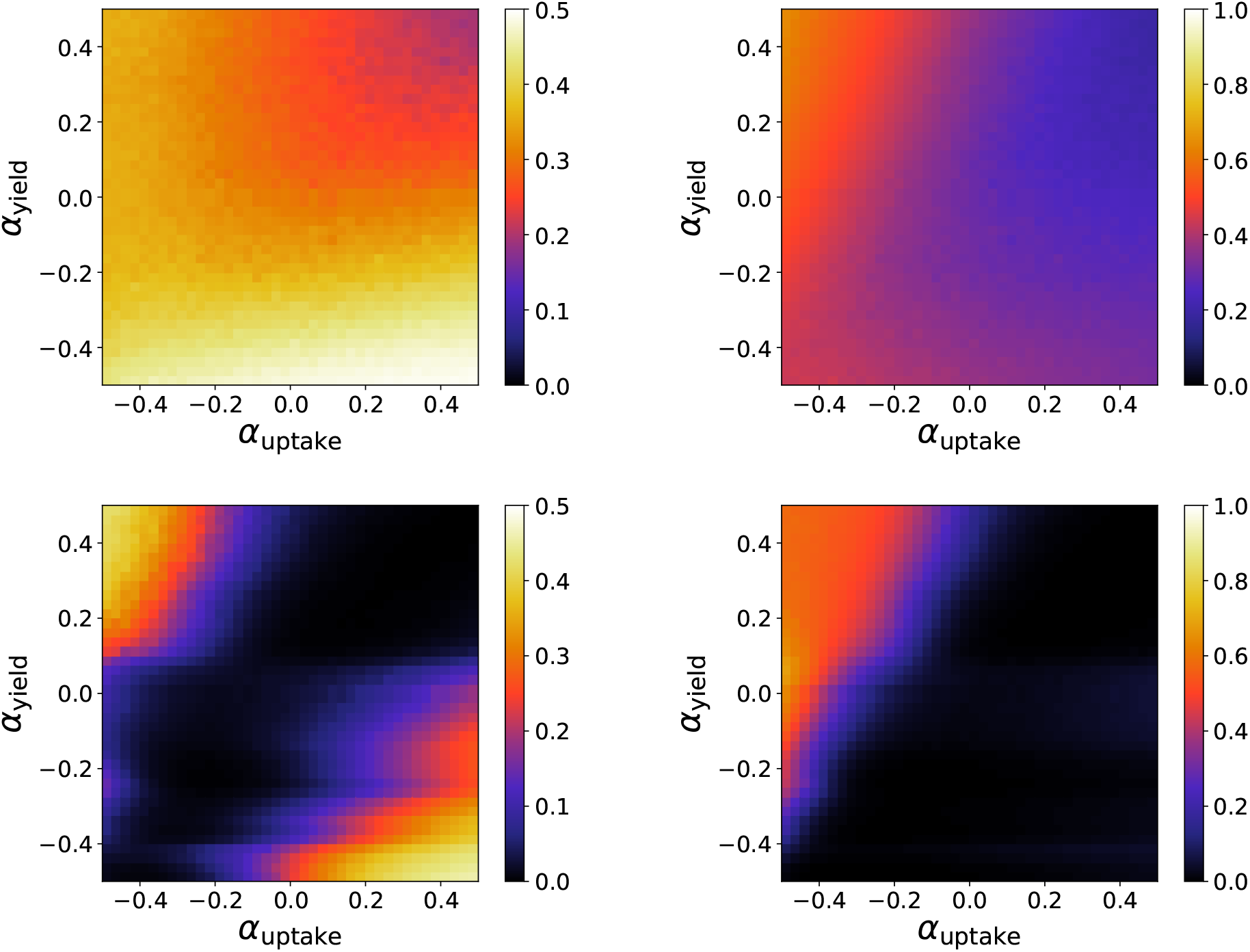
Specialization level for resource input rate Δ*R* = 1 (top panels) and Δ*R* = 200 (bottom panels). The left panels depict the specialization level in uptake and the right panels the specialization in yield. The parameters are *N* = 2, *ν* = 0.01, *A*_max_ = 10, *L* = 100, *D* = 0.2, Δ*t* = 1 and Δ*x* = 1. Each point consists of an average over 100000 time units, sampled every 50 time units, taken from 5 independent configurations.

Let us now scrutinize the response of the specialization to changes of the curvature parameter *α*_yield_. As all variables are somehow coupled through the definition of a surface in the 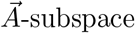, and in its turn the set of uptake rates {*A*^*k*^} are attached to the efficiencies {*η*^*k*^} through the *N* relations of the kind *A*^*k*^ × *η*^*k*^, *α*_yield_ is likewise expected to affect the dynamics as a whole. By making *α*_uptake_ constant, we observe that the rise of the curvature parameter *α*_yield_, which turns the tradeoff curves more concave, ensures reduced levels of specialization in the uptake of resources when resources are scarce (please, see left upper panel of Fig. 5)). Nevertheless, a more intricate situation materializes when Δ*R* is large, and now we may distinguish two different scenarios: over nearly all the domain of negative values of *α*_uptake_, specialization levels in resource uptake become a monotonic increasing function of *α*_yield_, while roughly over all the domain of positive *α*_uptake_-values the reversed condition is obtained, with specialization levels in resource uptake falling as *α*_yield_ grows.

In its turn, the dependence of specialization levels in high yield on the curvature parameter *α*_yield_ is less intricate. In general, we observe that the levels of specialization in high yield grow continuously with *α*_yield_, which is a quite impressive issue. Contrary to what would be expected under a scenario of single tradeoffs, specialization in high yield is here found when the relation between uptake rate and efficiency, *A*^*k*^ × *η*^*k*^, presents concave curvatures. Although the same pattern is found in the domain of positive values of *α*_uptake_, the levels of specialization in high yield are considerably smaller than when *α*_uptake_ takes negative values.

The above analysis is extended to the cases of *N* = 3 and *N* = 5 resource types. Roughly speaking, we can conclude that the general picture is very similar to that displayed for two resources (*N* = 2). However, we notice that when more resource types are incorporated into the dynamics the regime in which the specialization levels in resource uptake decrease with *α*_yield_ shrinks up to its disappearance, as already seen for *N* = 5. Therefore, when the number of resources *N* is not so small, the dependence of the specialization levels in resource uptake on *α*_yield_ is monotonic in the entire range of *α*_uptake_. Moreover, when Δ*R* is low, we also notice that specialization levels in resource uptake become less and less sensitive to variations on both curvature parameters. Another point to highlight is that specialization levels in high yield are considerably larger as more resource types are considered, as disclosed for *N* = 5, in which the levels of specialization are nearly the maximum over a broad domain of the diagram.

### 3.1 Microscopic analysis

For a more detailed comprehension of the phenomena captured in the previous plots, here we provide a microscopic analysis of the problem by exploring the spatial distribution of trait values. Fig. 8 displays snapshots of the spatial distribution of trait values and for different combinations of parameter values. In set of panels (a) the condition *α*_uptake_ = −0.3 and *α*_yield_ = 0.3 is addressed. For each resource type, there exists a group of individuals that achieve moderate to large uptake rate, but a substantial amount has uptake rate close to zero. By comparing the three snapshots we find out that nearly every individual achieves uptake rate very close to zero for at least one resource type. This accounts for the specialization levels unveiled in Fig. 6, which is around 0.3 for this particular combination of the curvature parameters. On the other hand, the same set of panels reveals that a considerable fraction of the individuals end up with high efficiencies, thus explaining the elevated specialization levels in high yield obtained, which is around 0.8, as demonstrated in Fig. 6. Keep in mind that in this case, specialization in high yield means that the ratio *η*^*k*^/(*A*^*k*^/*A*_*max*_) for a given resource is greater than a specified value *r*_*spec*_.

**Figure 6:**
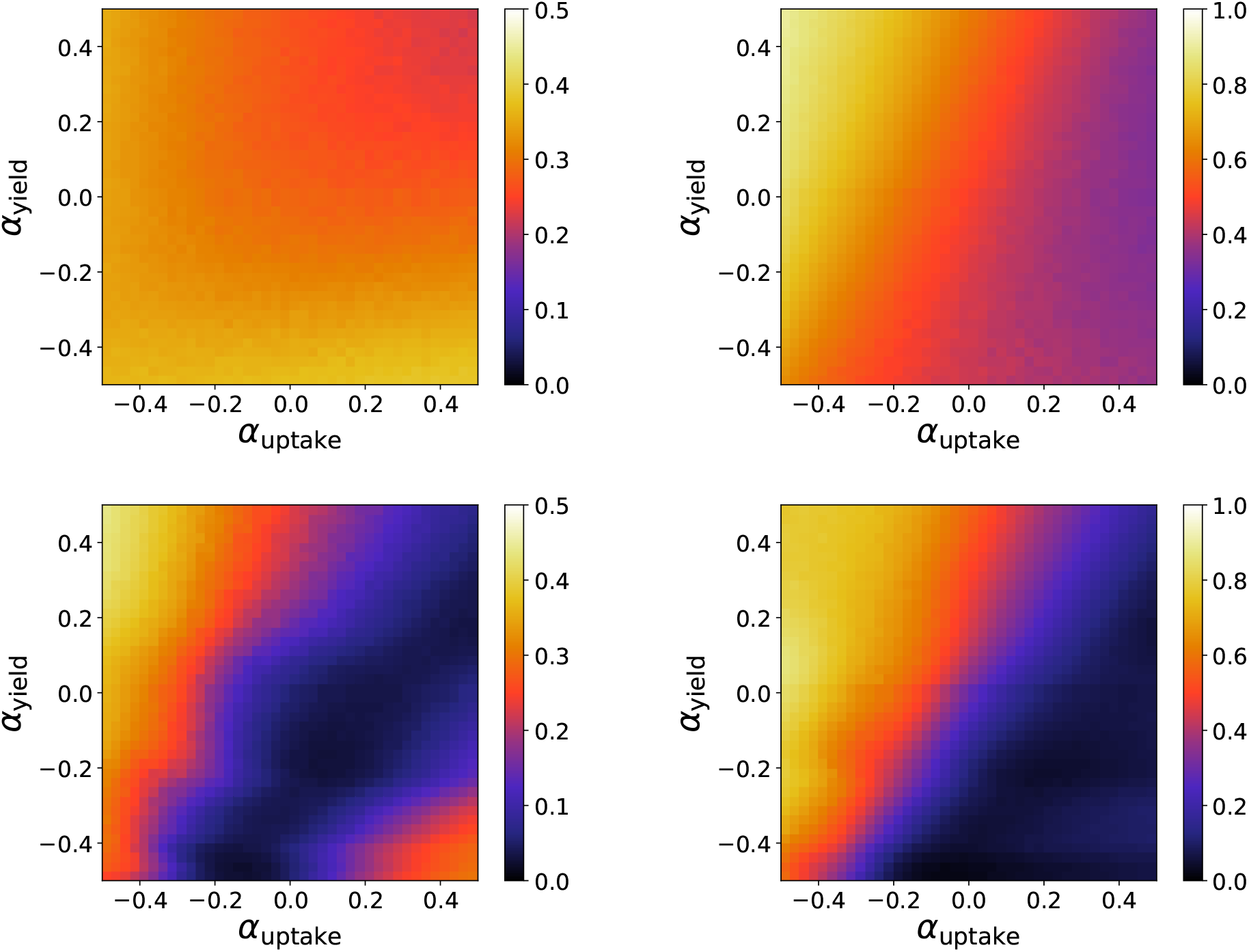
Specialization level for resource input rate Δ*R* = 1 (top panels) and Δ*R* = 200 (bottom panels). The left panels depict the specialization level in resource uptake and the right panels the specialization in high yield. The parameters are *N* = 3, *ν* = 0.01, *A*_max_ = 10, *L* = 100, *D* = 0.2, Δ*t* = 1 and Δ*x* = 1. Each point consists of an average over 100000 time units, sampled every 50 time units, taken from 5 independent configurations.

**Figure 7:**
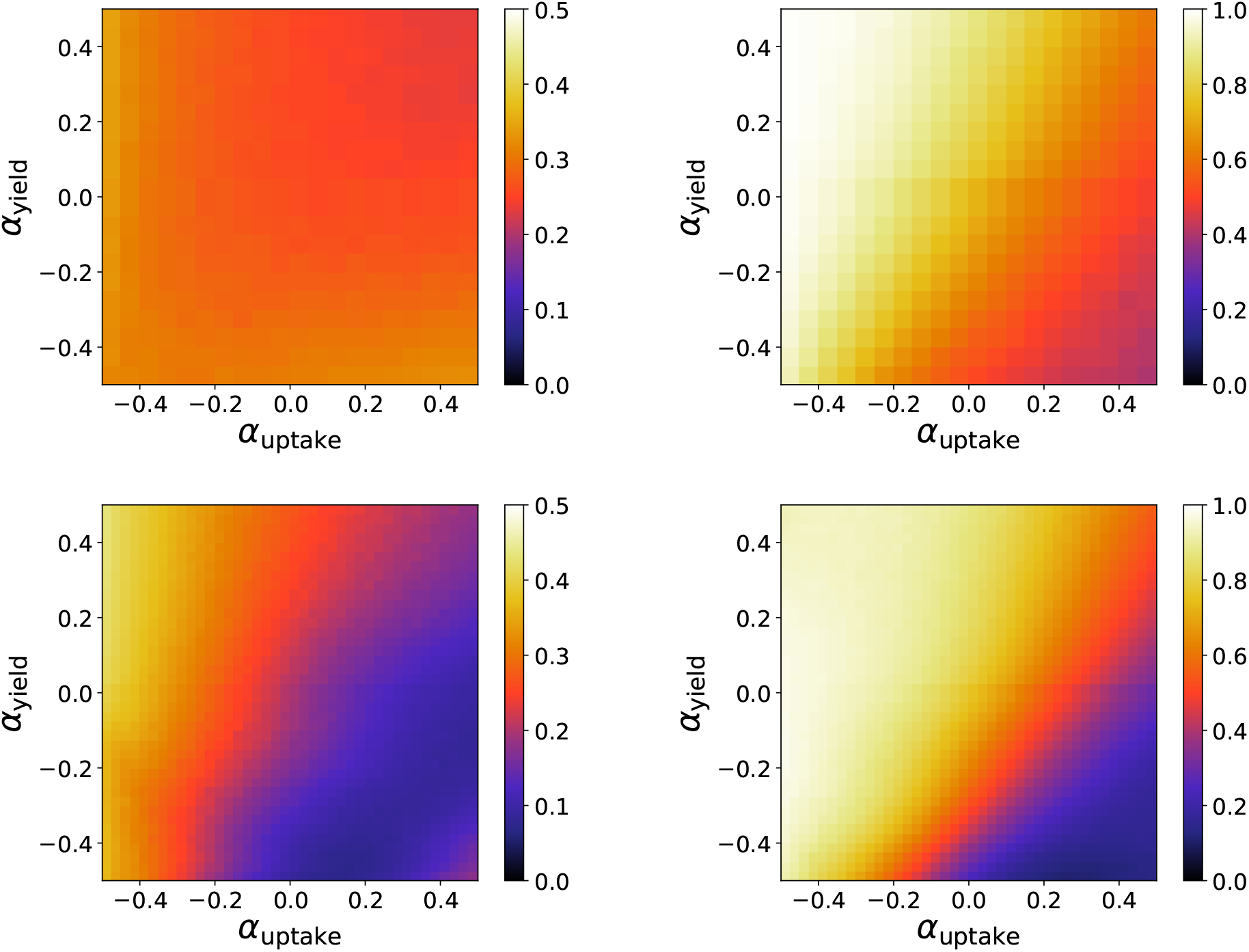
Specialization level for resource input rate Δ*R* = 1 (top panels) and Δ *R* = 200 (bottom panels). The left panels depict the specialization level in uptake and the right panels the specialization in yield. The parameters are *N* = 5, *ν* = 0.01, *A*_max_ = 10, *L* = 100, *D* = 0.2, Δ*t* = 1 and Δ*x* = 1. Each point consists of an average over 100000 time units, sampled every 50 time units, taken from 5 independent configurations.

**Figure 8:**
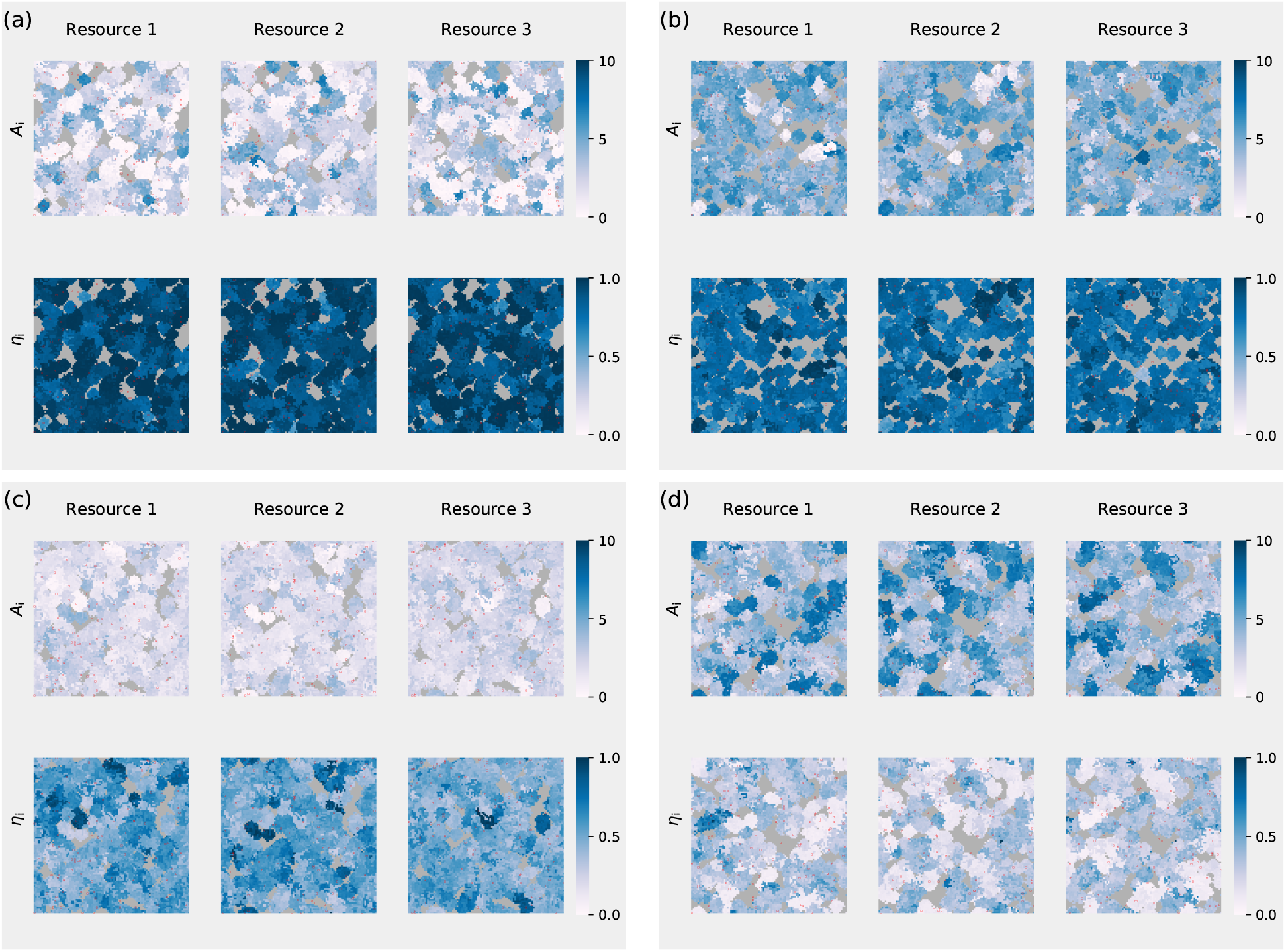
Snapshot of the spatial distribution of trait values. The number of resources is *N* = 3, and for each set the trait values of *A*^1^, *A*^2^ and *A*^3^; and *η*^1^, *η*^2^ and *η*^3^ are shown. The uptake and yield parameters are (a) *α* _uptake_ = − 0.3, *α*_yield_ = 0.3; (b) *α*_uptake_ = 0.3, *α*_yield_ = 0.3; (c) *α*_uptake_ = 0.3, *α*_yield_ = 0.3; (d) *α*_uptake_ = 0.3, *α*_yield_ = 0.3. For each set Each pixel represents a cell. The locations with resource input are highlighted with a red frame and the grey regions correspond to nonoccupied locations. The parameters are Δ*R* = 200, *N* = 3, *ν* = 0.01, *A*_max_ = 10, *L* = 100, *D* = 0.2, Δ*t* = 1 and Δ*x* = 1. The snapshot was obtained after 50000 time units.

In the set of panels (b), one has *α*_uptake_ = 0.3 and *α*_yield_ = 0.3. Note that now the distribution of uptake rates is much more uniform than found in panels (a). According to Fig. 6, in this scenario the specialization in both resource uptake and in high yield is very modest. The above scenario evinces that moderate to high uptake rates and efficiencies are not promoted at the expense of the other quantities. In panels (c), where *α*_uptake_ = *−*0.3 and *α*_yield_ = *−*0.3, once again the distribution of resource uptake rates looks flat, but contrary to the scenario in panels (b), the uptake rates are very low. The efficiencies are not so high as before, but still reach intermediate values.

Lastly, in set of Panels (d), in which *α*_uptake_ = 0.3 and *α*_yield_ = −0.3, one has a composition of individuals achieving low efficiencies and moderate uptake rates. As shown in Fig. 6, this combination of the curvature parameters leads to the loss of specialization in high yield and reduced levels of specialization in the uptake of resources.

In all the scenarios discussed above, the population is usually composed of relatively uniform superposing clusters of cells competing with each other for resources and space. These clusters allow relatively high diversity to be maintained in a global scale even though locally the diversity is low. When the resource influx is small the size of the clusters recedes and they become mostly disconnected. This can be observed in Fig. S5, provided in the Supplemental Information.

### 3.2 Random placement

The results presented until now do not allow us to disentagle the role played by the tradeoffs and the effect of structuring in promoting specialization. With this intent, we investigate a variant of the spatial model, here designated as random placement. The difference between the original and modified version is on the stage of reproduction. Instead of occupying a neighbouring site, one of the daughter cells is placed at a random empty location of the lattice, while the other one, as before, replaces the parental cell. If the lattice is fully populated, a daughter cell replaces a cell chosen at random in the lattice. In the original model, as kin cells tend to accumulate around a given location, kin selection is an effective selective pressure. The random placement model mitigates the contribution of kin selection to the evolutionary dynamics, but still preserves the underlying dynamics of resource distribution.

The dissolution of any form of spatial pattern formation among individuals is clearly demonstrated in Fig. S6 (Supplemental Information). The patterns of trait-values distributions as exhibited in Fig. 8 are now substituted by homogeneous distributions over the lattice. Under this scheme, specialization in resource uptake is not expected, as corroborated in Fig. 9. On the other hand, specialization in high yield will be possible whenever the efficiencies are high and the uptake rates are low. This situation is observed in panels (a) and (c), and quantitatively established in the right panel of Fig. 9. So, opposed to the drastic effects of random placement on the specialization levels in resource uptake, those effects are much less influential in shaping the specialization levels in high yield and the outcomes are similar to those seen for the original model. The specialized spatial domains observed in the original model are an essential mechanism of promoting diversity of strategies regarding the uptake of resources.

**Figure 9:**
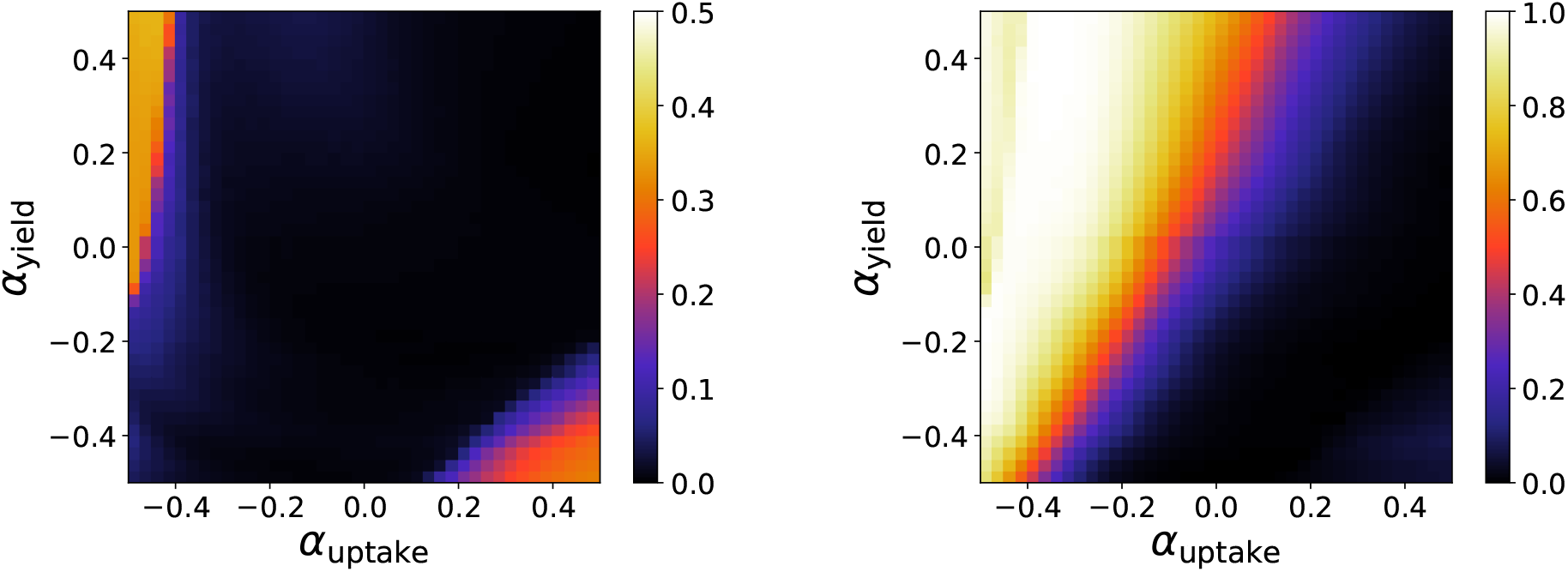
Specialization level for resource input rate Δ *R* = 200 in a random placement population. The left panel depicts the specialization level in uptake and the right panel the specialization in yield. The parameters are *N* = 3, *ν* = 0.01, *A*_max_ = 10, *L* = 100, *D* = 0.2, Δ*t* = 1 and Δ*x* = 1. Each point consists of an average over 100000 time units, sampled every 50 time units, taken from 5 independent configurations.

## 4 Conclusions

Tradeoff relationships are a critical factor in controlling the degree of ecological specialization in natural populations (Michod et al. 2006). From an evolutionary perspective, this issue has been largely debated within the context of germ-soma differentiation (Michod et al. 2006; Leslie et al. 2017), and also discussed within the context of the evolution of multicellularity (Gavrilets 2010; Ispolatov et al. 2012; Amado and Campos 2017; Amado et al. 2018). A central feature about the tradeoffs is their structure (Jessup and Bohannan 2008; Saeki et al. 2014). Within an ecological viewpoint, the role of the shape of tradeoffs has been related to the emergence of specialists and/or generalists individuals (Egas et al. 2004; Guillaume and Otto 2012), to the levels of biodiversity within communities (Farahpour et al. 2018), and even to their responses to ecological competition (Maharjan et al. 2013). The most diverse forms of tradeoff have been reported, particularly some of those describing relationships between life-history traits are well established in the literature (Jessup and Bohannan 2008; Saeki et al. 2014).

Nevertheless, the usual approach to describe the tradeoff as single curves relating any two traits that are implicitly subject to biological and biophysical constraints is no longer suitable in case those traits are mediated by other mechanisms or traits. Therefore, a proper description of such cases requires a multivariate analysis to properly address the complex structure of trait spaces (Edwards and Stachowicz 2010; Edwards et al. 2011). Here, we have assumed the existence of multivariate tradeoff patterns to address the development of ecological specialization. The traits comprising the trait space are associated with metabolic properties of the individuals, more specifically, their resource uptake rates and corresponding efficiencies. The current approach allows us to observe how the structure of the multidimensional tradeoffs regulates the levels of specialization in a scenario where those traits are intertwined and compare how the new predictions differ from classical ecological studies of two traits. The levels of ecological specialization are studied in a scenario of a harsh environment, e.g. low resource influx rate, as well as in a scenario where resources abound. Moreover, a thorough survey of how the shape of the tradeoff relationships affects specialization is accomplished.

Our results show that the adaptive responses of the population are not uniquely determined by the shape of the tradeoffs, but also influenced by environmental variables, which is in accordance with the findings of Jessup and Bohannan (Jessup and Bohannan 2008). For instance, while at large resource input the population size declines with *α*_yield_, at small resource input the population grows with *α*_yield_. This fact is even more pronounced for *N* = 2 and *N* = 3. Regarding the measurements for the specialization levels, it is possible to state that they are normally enhanced upon harsh conditions. The pattern exhibited by the specialization levels in high yield in terms of both *α*_uptake_ and *α*_yield_ is qualitatively not much changed when the resource input and number of resources *N* are altered. In general, we find that the specialization levels in high yield are likely to increase as *α*_yield_ rises, which is a quite remarkable event, pointing out that any inference about the adaptive responses in populations subjected to multidimensional tradeoffs can not be concluded using the way of thinking of the classical ecological models of two-traits.

Quite intricate scenarios are exhibited by the specialization levels in resource uptake when the resource influx and/or the number of resources *N* are modified. Especially for *N* = 2 and *N* = 3, we observe that the specialization levels in resource uptake decline as *α*_uptake_ increases in the domain of negative *α*_yield_, but increases or is maximized at intermediate *α*_uptake_ in the domain of positive *α*_yield_. In a nutshell, the convolution of different tradeoffs can effectively lead to quite complex scenarios, as observed here.

Finally, the role of structuring on promoting specialization is examined. We proposed an alternative model in which the effect of kin selection can be completely attenuated by allowing daughter cells to occupy randomly chosen sites of the lattice instead of keeping constrained to the neighborhood of the parental cells. While the specialization levels in high yield in terms of *α*_uptake_ and *α*_yield_ resemble those found for the structured model, specialization in resource uptake, in essence, fades away. This results clearly prove that in such scenarios the generalist strategy provides the best selective alternative, with the resource uptake rates assuming small or moderate values.

The model here proposed has the potential for generalization along several lines (check section **??**of Supplemental Information for some possible directions). Besides the results obtained and discussed in this work, we introduced a general and flexible model that can itself be used to parameterize complex tradeoff surfaces in other works.

## Acknowledgments

PRAC acknowledges financial support from Conselho Nacional de Desenvolvimento Científico e Tecnológico (CNPq) under Projects No. 302569/2018-9 and 406594/2018-0. AA has a fellowship from the program PNPD sponsored by CoordenaÇão de AperfeiÇoamento de Pessoal de Nível Superior (CAPES).

## Supplemental Information

### A Summary of the parameters

#### Lattice parameters and quantities

*L*: linear size of the lattice
{*i, j*}: lattice site indexed by *i* and *j*
Δ*x*: lattice spacing (distance between neighboring sites)
Δ*t*: time step

#### Resource parameters and quantities

*N*: number of resources
*n*_loc_: number of resource input sites
Δ*R*: total resource inflow rate in the lattice
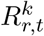: amount of resource *k* at site *r* and time *t*
*D*: resource diffusion constant

#### Cell parameters and quantities

*A*^*k*^: uptake rate
*A*_max_: maximum achievable uptake rate
*η*^*k*^: efficiency of resource processing
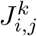: rate of resource acquisition
Δ*E*_*i,j*_: amount of energy acquired by an individual in a time step
*E*_split_: energy threshold for reproduction
*ν*: individual death rate
*α*_uptake_: curvature parameter of uptake tradeoff
*α*_yield_: curvature parameter of uptake-yield tradeoff

Note: Throughout the text, superscripts are added to quantities to specifiy the resource, while subscripts are used to indicate either the lattice position and time (in the case of resource related quantities) or the individual cell (in the case of cell related quatities). Typically, an index *k* is used for the resource, an index *r* for the position, an index *t* for the time and an index *i* for the individual cell. As an illustration, 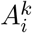 is the uptake rate of cell *i* for resource *k* and 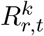 is the amount of resource *k* present at the lattice site *r* and time instant *t*.

### B Single tradeoff

**Figure S1:**
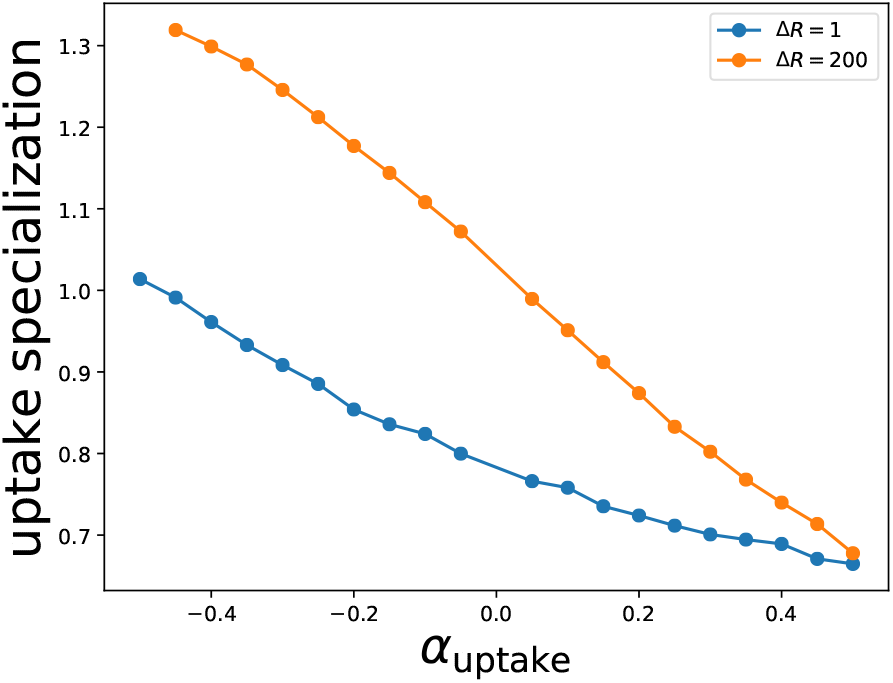
Specialization level in uptake for resource input Δ*R* = 1 and Δ*R* = 200 when only the tradeoff between uptake rates is considered. The efficiencies *η*^*k*^ have been kept fixed at 0.5, independently of the value of *A*^*k*^. One can see that when the tradeoff becomes less convex and more concave the specialization level falls monotonically, in accordance with the classic expectation. The parameters are *N* = 3, *ν* = 0.01, *A*_max_ = 10, *L* = 100, *D* = 0.2, Δ*t* = 1 and Δ*x* = 1. Each point consists of an average over 100000 time units, sampled every 50 time units, taken from 50 independent configurations.

### C Lattice occupation

**Figure S2:**
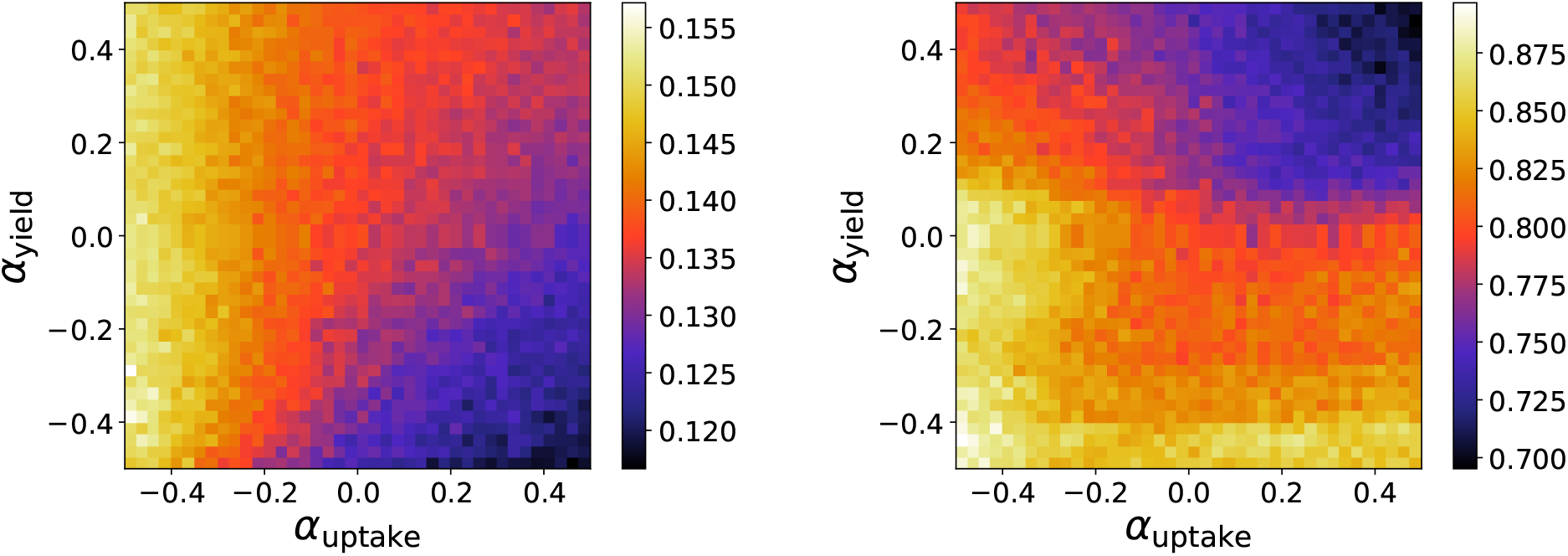
Lattice occupation for resource input rate Δ*R* = 1 (left panel) and Δ*R* = 200 (right panel). The parameters are *N* = 2, *ν* = 0.01, *A*_max_ = 10, *L* = 100, *D* = 0.2, Δ*t* = 1 and Δ*x* = 1. Each point consists of an average over 100000 time units, sampled every 50 time units, taken from 5 independent configurations.

**Figure S3:**
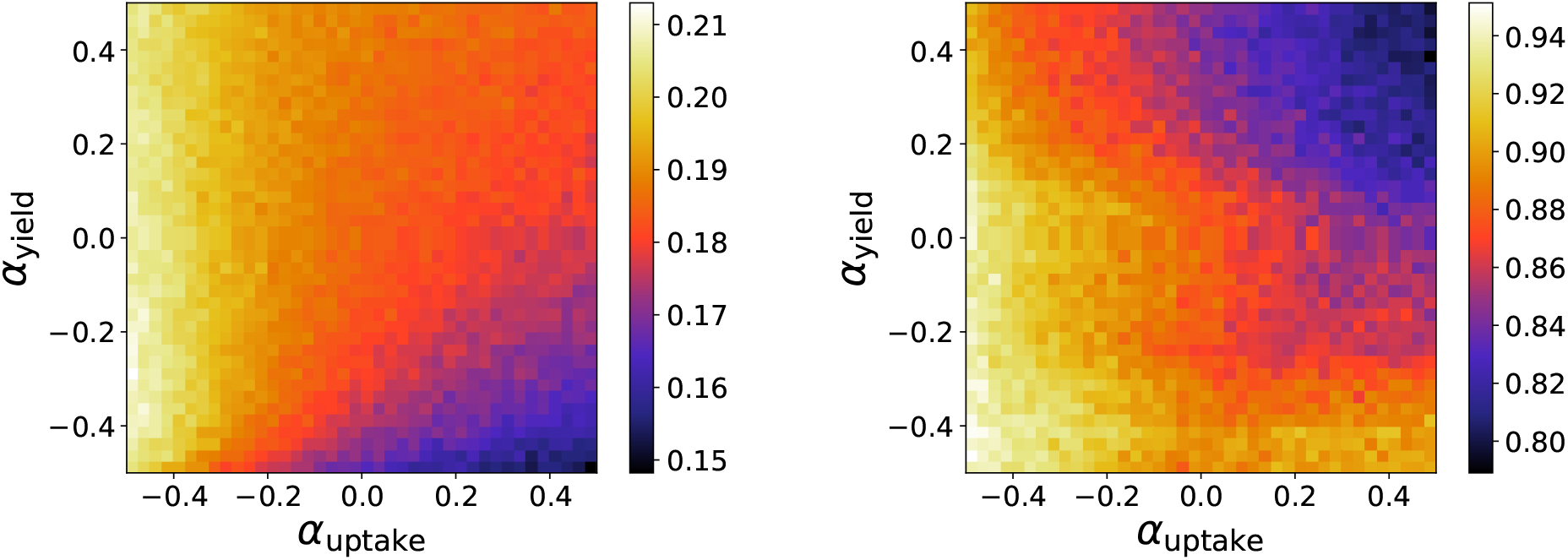
Lattice occupation for resource input rate Δ*R* = 1 (left panel) and Δ*R* = 200 (right panel). The parameters are *N* = 3, *ν* = 0.01, *A*_max_ = 10, *L* = 100, *D* = 0.2, Δ*t* = 1 and Δ*x* = 1. Each point consists of an average over 100000 time units, sampled every 50 time units, taken from 5 independent configurations.

**Figure S4:**
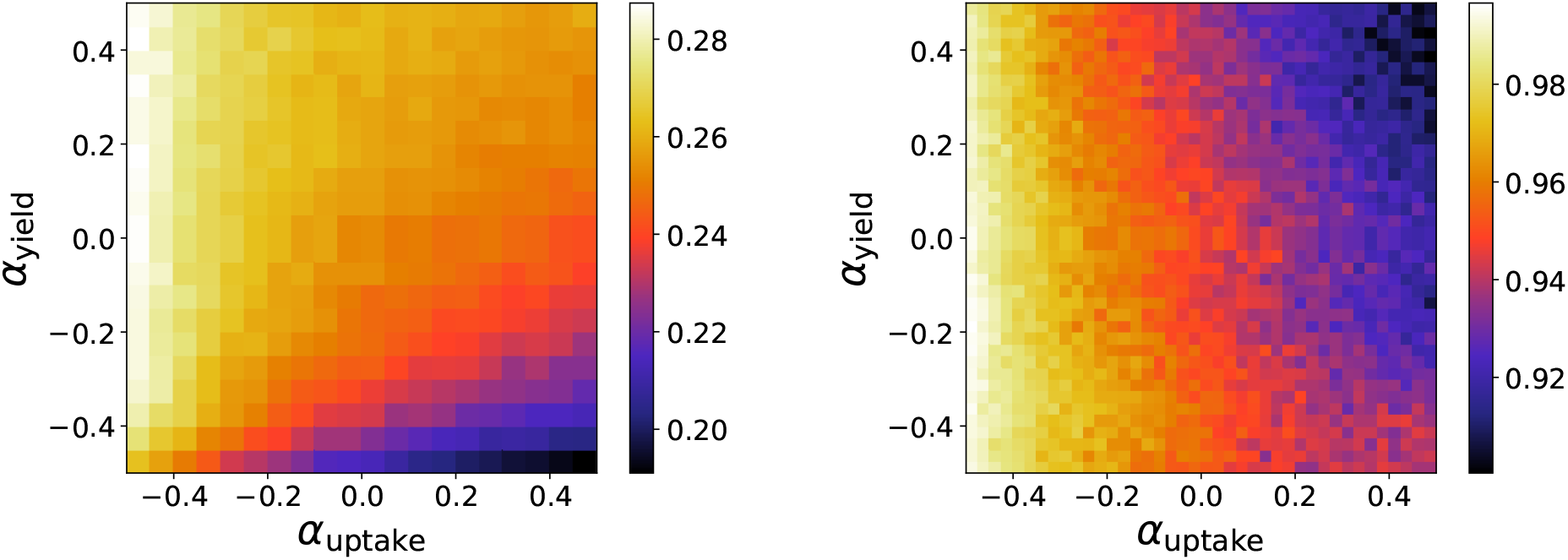
Lattice occupation for resource input rate Δ*R* = 1 (left panel) and Δ*R* = 200 (right panel). The parameters are *N* = 5, *ν* = 0.01, *A*_max_ = 10, *L* = 100, *D* = 0.2, Δ*t* = 1 and Δ*x* = 1. Each point consists of an average over 100000 time units, sampled every 50 time units, taken from 5 independent configurations.

### D Spatial snapshots

**Figure S5:**
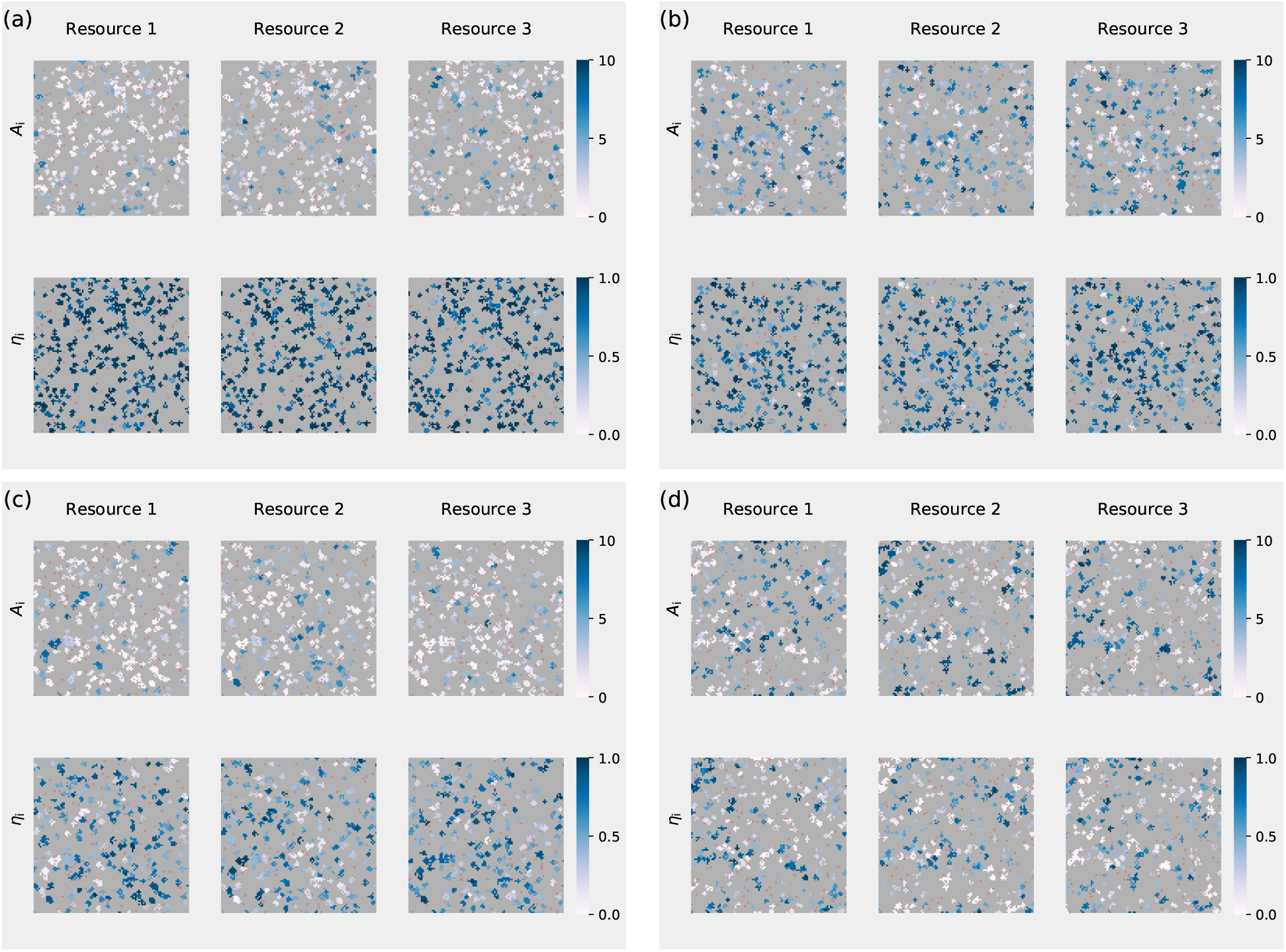
Snapshots of the spatial distribution of trait-values. The uptake and yield parameters are (a) *α*_uptake_ = −0.3, *α*_yield_ = 0.3; (b) *α*_uptake_ = 0.3, *α*_yield_ = 0.3; (c) *α*_uptake_ = −0.3, *α*_yield_ = −0.3; (d) *α*_uptake_ = 0.3, *α*_yield_ = −0.3. Each pixel represents a cell. The locations with resource input are highlighted with a red frame and the grey areas correspond to nonoccupied locations. The parameters are Δ*R* = 1, *N* = 3, *ν* = 0.01, *A*_max_ = 10, *L* = 100, *D* = 0.2, Δ*t* = 1 and Δ*x* = 1. The snapshot was obtained after 50000 time units.

### E Random placement

**Figure S6:**
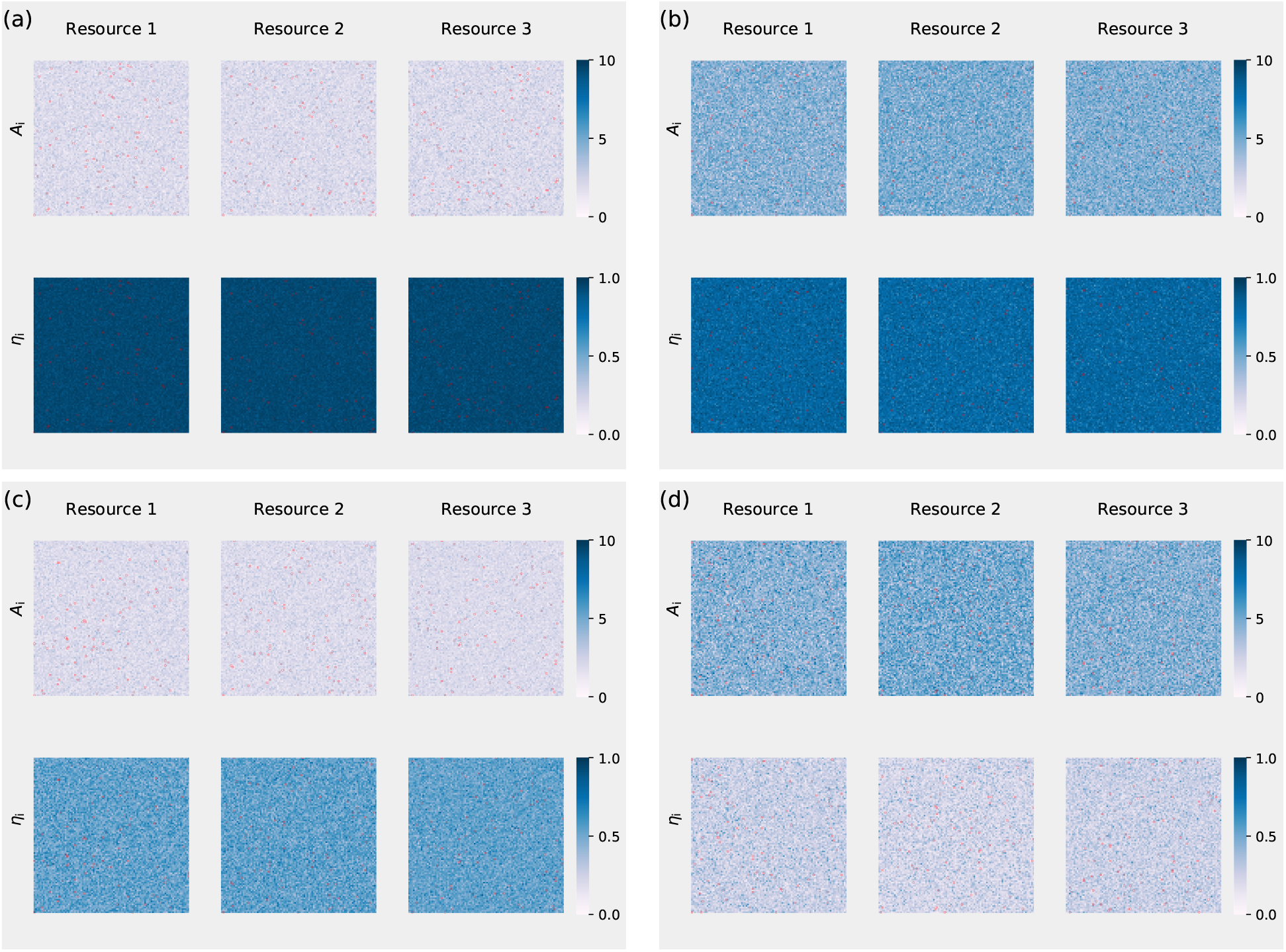
Snapshots of the spatial distribution of trait-values for the random placement model. The uptake and yield parameters are (a) *α*_uptake_ = −0.3, *α*_yield_ = 0.3; (b) *α*_uptake_ = 0.3, *α*_yield_ = 0.3; (c) *α*_uptake_ = −0.3, *α*_yield_ = −0.3; (d) *α*_uptake_ = 0.3, *α*_yield_ = −0.3. Each pixel represents a cell. The locations with resource input are highlighted with a red frame and the grey regions correspond to nonoccupied locations and the grey areas correspond to nonoccupied locations. The parameters are Δ*R* = 200, *N* = 3, *ν* = 0.01, *A*_max_ = 10, *L* = 100, *D* = 0.2, Δ*t* = 1 and Δ*x* = 1. The snapshot was obtained after 50000 time units.

### F Generalizations

This work can be generalized in a number of ways. The simplest generalizations are probably releasing the requirement of symmetry among resources. One way to do this is to consider a different 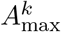 for each resource. Another possibility is to assign different curvatures *α*^*j*×*k*^ to each tradeoff *j × k*. Figs. S7 shows example tradeoff surfaces for which these assumptions have been released. To implement these generalizations one only has to modify eq. (7) (main text) so that it takes the form

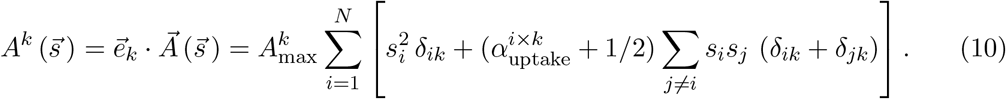

**Figure S7:**
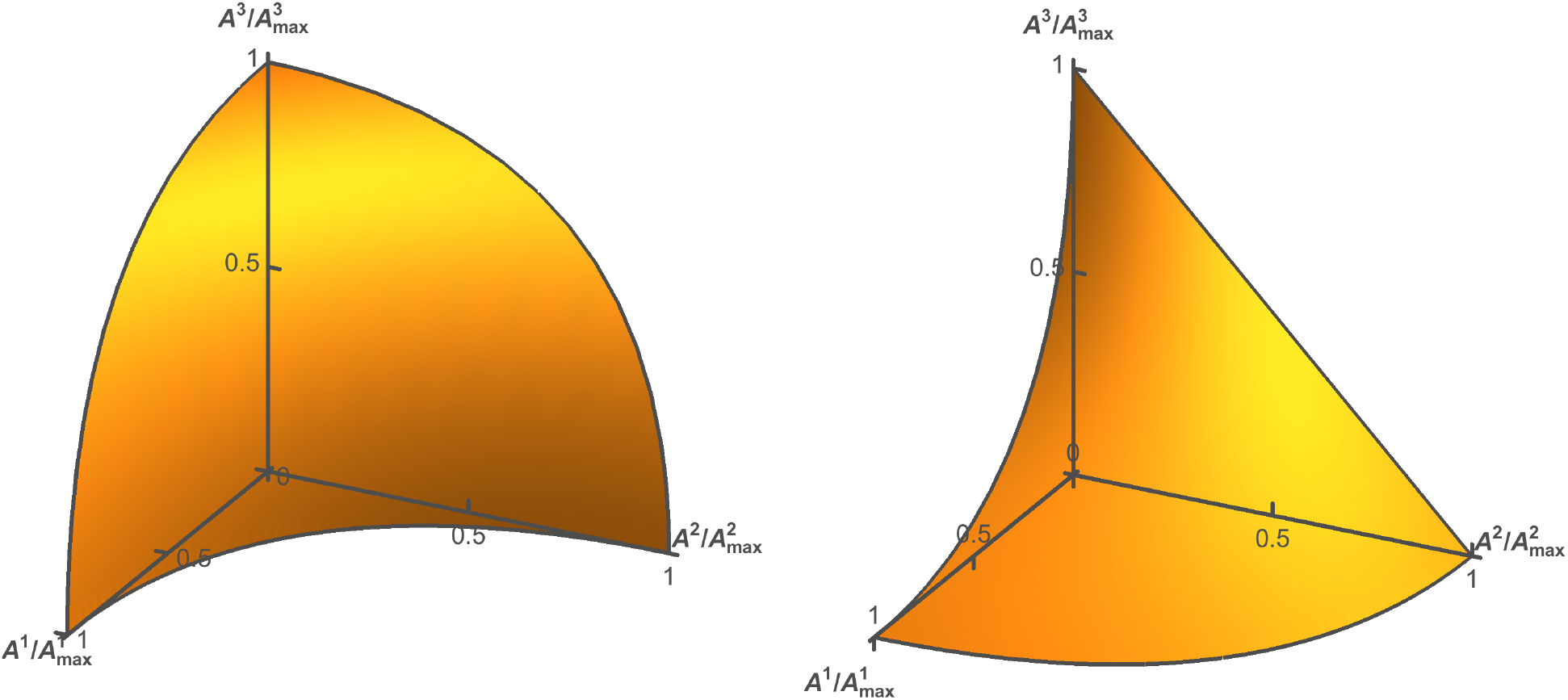
Examples of generalized tradeoff surfaces for independent *α^j×k^* for each tradeoff *j × k*. Left panel: *α*^1 *×*2^ = −0.5, *α*^2 *×*3^ = 0.5 and *α*^1 × 3^ = 0.5. Right panel: *α*^1 × 2^ = 0.5, *α*^2 × 3^ = 0 and *α* ^1 × 3^ = −0.5.

